# DNARecords: An extensible sparse format for petabyte scale genomics analysis

**DOI:** 10.1101/2022.08.13.503863

**Authors:** Andres Manas, Lucas Seninge, Atray Dixit

## Abstract

Recent growth in population scale sequencing initiatives involve both cohort scale and proportion of genome surveyed, with a transition from genotyping arrays to broader genome sequencing approaches. The resulting datasets can be challenging to analyze. Here we introduce DNARecords a novel sparse-compatible format for large scale genetic data. The structure enables integration of complex data types such as medical images and drug structures towards the development of machine learning methods to predict disease risk and drug response. We demonstrate its speed and memory advantages for various genetics analyses. These performance advantages will become more pronounced as it becomes feasible to analyze variants of lower population allele frequencies. Finally, we provide an open-source software plugin, built on top of Hail, to allow researchers to write and read such records as well as a set of examples for how to use them.

## Introduction

Modern population scale sequencing projects involve sequencing hundreds of thousands to millions of individuals in which genetics data can be correlated with a broad assortment of healthcare data. The number of variants that can be analyzed has grown substantially from initial efforts with genotyping arrays to recent releases of whole genome sequencing (WGS) data from both the UK Biobank [1] and the All of Us initiative [2]. As the size of these datasets has grown, new methods have been developed to facilitate their analysis. Tools such as Hail [3] and REGENIE [4] are useful for performing genetic analysis, such as fixed or random effects GWAS efficiently on large patient cohorts in a distributed framework that can be deployed on the cloud. Other frameworks have been developed to enable GPU based operations on genetics datasets [5], including QTL analysis [6], for improved speed, but are oriented for variant-by-variant analysis.

One goal of large population sequencing initiatives is the generation of novel precision medicine solutions to help stratify patients with respect to disease risk (including polygenic risk scores) and therapy selection. Currently, genetics informed clinical predictors are approaching clinical utility for certain complex diseases, including predictors of cardiac risk [7]. Healthcare outcomes are a result of genetic (G), environmental (E), sociodemographic (S), and random effects. There are additional clinical variables that can be informative. Most current approaches for predicting healthcare outcomes using genetic data take into account simple numerical covariates (S and E) such as age or lipid levels. A modeling approach that can take into account other important, but harder to model data types (such as medical images or drug structure) is likely to result in improved predictors for critical outcomes such as hospitalization risk [8], [9]. The integration of these complex non-tabular datatypes is made possible by modern deep learning methods (including forms of convolutional neural network architectures), which operate on datasets oriented in a sample-wise fashion. Lastly, there is significant evidence that there is a small, but significant non-additive component to the heritability of complex human traits [10], [11]. Modeling these nonlinearities is facilitated by data structures organized by sample. As such, there is a need for analysis frameworks that restructure genetics datasets in a machine learning compatible format (i.e. the transpose of traditional genetics data storage formats).

Here we introduce DNARecords, a new genetics data format that has significant advantages for speed and novel machine learning applications. The format leverages sparsity in genetics datasets for faster computation and can be stored in a sample-wise manner for integrations of complex data types. Moreover, by working on top of existing frameworks including Tensorflow’s TFRecords and Apache Parquet, datasets can be distributed in the cloud for analysis as well as across GPUs or TPUs [12].

## Results and Discussion

DNARecords stores genetic data in both sample-wise and variant-wise data structures in which sparsity is leveraged (**Figure 1A**). Specifically, entries where the genotypes are homozygous for the reference allele (or relatedly when genotype dosage is below a certain threshold) can be stored implicitly. We show that when using this format to perform genetics analysis like GWAS there are negligible losses in sensitivity or specificity to detect significant variants when the dosage threshold is less than 0.1 (**Figure 1C**). The performance advantages of DNARecords in memory footprint for several example datasets are described below (**Table 1**). These advantages become pronounced with larger WES and WGS datasets, in which progressively rare variants with lower population allele frequencies are considered for analysis.

**Table 1:** Footprint comparison of DNARecords to other genetics datatypes as a function of genotyping assay.

| Dataset | Number of samples | Number of variants | Average allele frequency | HWE estimated sparsity factor<sup>3</sup> |
| --- | --- | --- | --- | --- |
| Genotyping array<sup>4</sup> | 3,000 | 693,158 | 0.1434 | 3.8 |
| Imputed genotypes AF>0.001<sup>3</sup> | 490,030 | 13,000,000 | 0.081 | 6.4 |
| Imputed genotypes<sup>5</sup> | 490,030 | 100,000,000 | 0.021 | 24 |
| WGS<sup>4</sup> | 76,156 | 759,302,267 | 0.0043 | 116 |
| WES<sup>6</sup> | 125,748 | 17,201,296 | 0.0016 | 312 |

**Figure 1:**
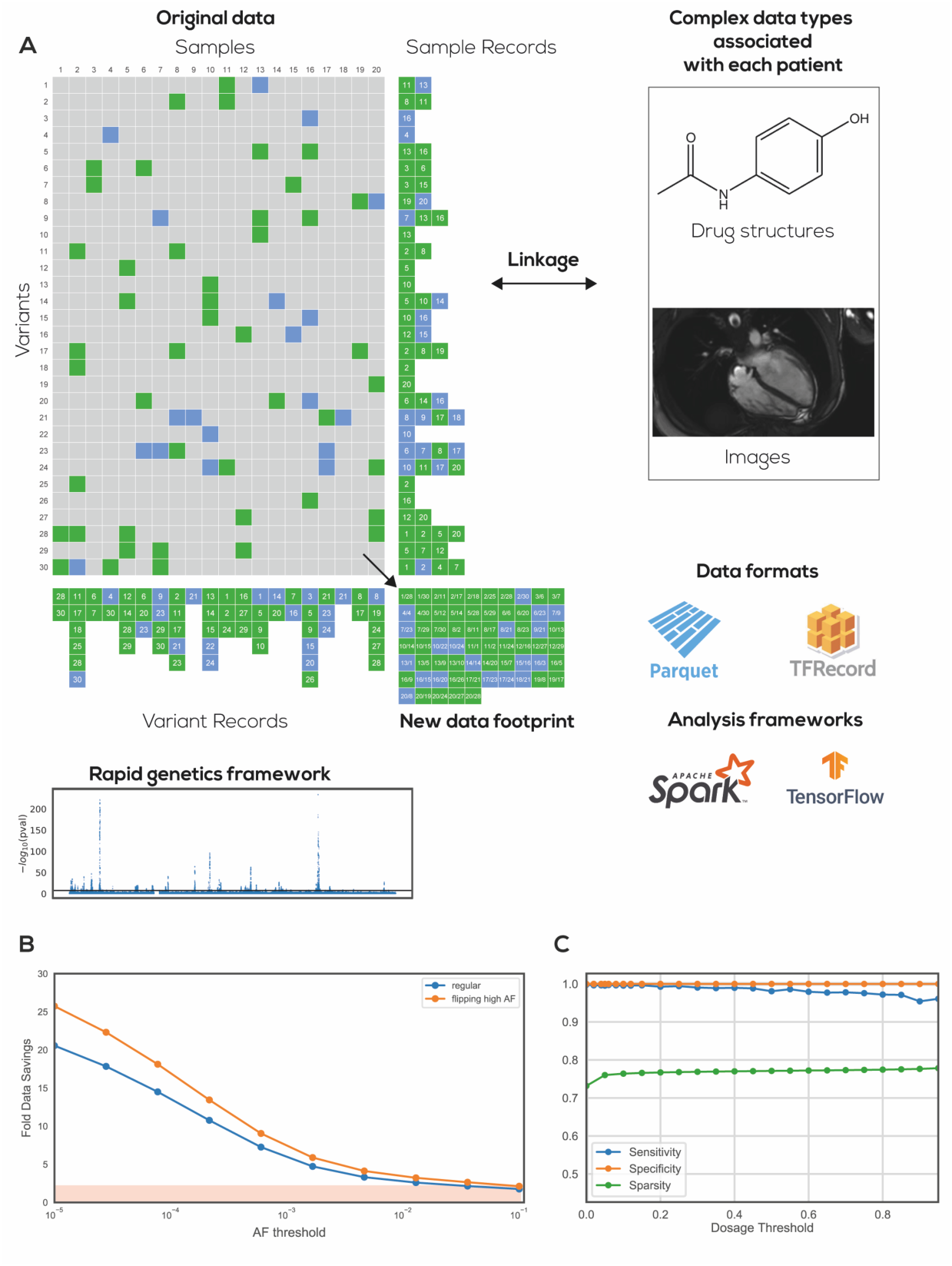
(**A**) DNARecords data format stores variant data separately in sparse sample oriented and variant oriented files to allow for efficient parallelism on standard genetics operations as well as new machine learning approaches. Sample-wise records can be linked to complex patient specific datatypes such as graph-encoded chemical structures and medical images (**B**) Data savings associated with thresholding variants above different allele frequency cutoffs. (**C**) Relationship between increasing sparsity (green) by adjusting the threshold for dosage and sensitivity (blue) /specificity (green) in a GWAS for height in the UK Biobank on chromosome 22.

These dataset size advantages are propagated into faster analysis for operations like GWAS and PCA (see **Table 2**).

**Table 2:** CPU hours. Speed comparisons for basic genetics operations for 2.5M variant dataset with the 1,000 genomes dataset. Numbers extrapolated based on the UK Biobank dataset.

| Method | GWAS | PCA | 2-layer model with images |
| --- | --- | --- | --- |
| Hail | 1.11 | 23.73 | N/A |
| DNAREcords (+1 P100 GPU) | 0.55 | 1.16 | 1.15 |

## Methods

The basic procedures for data generation are described here: https://dnarecords.readthedocs.io/. A python package for generating DNARecords for variant-wise analysis or sample wise machine learning is available here: https://pypi.org/project/dnarecords/. Open source code is available in Github here: https://github.com/amanas/dnarecords. DNARecords datasets for the 1kg dataset are hosted in Google Cloud storage (in both TFRecords and Parquet format) here: gs://dnarecords/1kg

## Acknowledgements

This work was supported by an NIH NHGRI 2R44HG010445-02.

## Author contributions

A.M. developed the data pipeline to transform standard genetics formats into DNARecords. L.S., A.M., and A.D. performed computational analysis. A.D. drafted the original manuscript. A.M., L.S., and A.D. revised the manuscript.

## Competing interests

A.D. is an equity holder in Coral Genomics, Inc.

## Footnotes

3 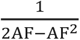

4 based on 1,000 genomes

5 based on UK Biobank

6 based on gnomAD

## Notes

https://dnarecords.readthedocs.io/

